# Cellular adaptation and the importance of the purine biosynthesis pathway during biofilm formation in Gram-positive pathogens

**DOI:** 10.1101/2020.12.11.422287

**Authors:** Martin Gélinas, Léa Museau, Arielle Milot, Pascale B. Beauregard

## Abstract

Bacterial biofilms are involved in chronic infections and confer 10 to 1000 times more resistance to antibiotics, leading to treatment failure and complications. When transitioning from a planktonic lifestyle to biofilms, certain Gram-positive bacteria are likely to modulate several cellular pathways including central carbon metabolism, primary biosynthesis pathways and production of secondary metabolites. These metabolic adaptations might play a crucial role in biofilm formation by Gram-positive pathogens such as *Staphylococcus aureus* and *Enterococcus faecalis*. Here, we performed a transcriptomic approach to identify cellular pathways that might be similarly regulated during biofilm formation in these bacteria. Different strains and biofilm-inducing media were used to identify a set of regulated genes that are common and independent of the environment or accessory genomes analysed. The gene set enrichment analysis of the transcriptome of four different strains of Gram-positive bacteria identified biosynthesis of secondary metabolites, biosynthesis of antibiotics and purine biosynthesis as three commonly upregulated pathways in biofilm. Our approach did not highlight downregulated pathways during biofilm formation that were common to *S. aureus* and *E. faecalis*. Of the three upregulated pathways, the *de novo* IMP biosynthesis pathway constitutes a promising target of cellular adaptation during biofilm formation. Gene deletions in this pathway, particularly *purN, purL, purQ, purH* and *purM* significantly impaired biofilm formation of *S. aureus*.

**Importance:** Biofilms are often involved in nosocomial infections and can cause serious chronic infections if not treated properly. Current anti-biofilm strategies rely on antibiotic usage, but they have a limited impact because of the biofilm’s intrinsic resistance to drugs. Metabolism remodelling likely plays a central role during biofilm formation, but it is still unclear if these cellular adaptations are shared between strains and species. Using comparative transcriptomics of different strains of *Staphylococcus aureus* and *Enterococcus faecalis*, we identified a core of commonly regulated genes during biofilm formation. Interestingly, we observed that the *de novo* IMP biosynthesis was systematically upregulated during biofilm formation. This pathway could constitute an interesting new anti-biofilm target to increase the host spectrum, drug efficiency and prevent resistance evolution. These results are also relevant to a better understanding of biofilm physiology.

## Introduction

Biofilms are ubiquitous in the biosphere, since approximately 80% of all bacteria adopt this lifestyle (1). These structures are formed by multicellular communities that adhere to a surface or an interface and are embedded in a matrix of extracellular polymeric substances (EPS) (2, 3). The intrinsic properties of the matrix confer them the ability to adapt to their environment and resist hostile conditions (2). Biofilms are known to cause significant problems in different aspects of human activity such as the food industry and the industrial and clinical sectors (4–7).

According to the National Institutes of Health (NIH), up to 65% of nosocomial microbial infections and 80% of chronic infections involve bacterial biofilm (8). In hospitals, patients with biomedical implants such as prostheses are particularly at risk of developing severe complications related to biofilm-mediated infection during or after their surgery. These complications increase the morbidity and mortality rates of patients, as well as the duration and costs of hospitalization (9, 10). The most common pathogens isolated from those types of infection are *Staphylococcus aureus*, followed by *Enterococcus* spp., in approximately 36% and 19% of cases respectively (11, 12). Intrinsic properties of their biofilms provide 10 to 1000 times more resistance to antibiotics than the minimal inhibitory concentration (MIC) required to kill their planktonic counterparts (10, 13–16). Therefore, biomedical implant infections caused by biofilm can be particularly difficult to treat because they are often unresponsive to antibiotic lock therapy (ALT), the currently recommended clinical practice to prevent and treat catheter-related infections (17).

*S. aureus* is a Gram-positive bacterium that naturally composes up to 60% of the population, but it can also become an opportunistic pathogen, causing a wide range of infections acquired in hospital or community environment (18). *S. aureus* biofilms often contain a matrix mainly composed of the polysaccharide intercellular adhesin (PIA), an important component of cell-cell adhesion (19). PIA production is regulated by the *icaADBC* operon (20) and is induced by high concentration of glucose and salt in the medium (21, 22). However, while most *S. aureus* strains possess the *ica* operon, some of them produce a PIA-independent biofilm (22). In these biofilms, the matrix is mostly composed of extracellular DNA (eDNA) and large proteins that form fibrils for intercellular adhesion (23). *S. aureus* resistant to methicillin (SARM) and *S. aureus* sensitive to methicillin (SASM) are more likely to produce PIA-independent and PIA-dependant biofilm, respectively (24, 25). Although *S. aureus* biofilms are well studied, certain properties of PIA-dependant and PIA-independent biofilms are still poorly understood (26).

*Enterococcus faecalis* is a Gram-positive bacteria primarily found as a commensal organism that colonizes in low abundance our gastrointestinal tract (27). In hospitals, *E. faecalis* can become an opportunistic pathogen causing several potentially lethal infections in susceptible patients (28). *E. faecalis* possesses a high level of tolerance towards environmental stress and antibiotic treatments (29–32), probably due to its ability to form a biofilm (33). *E. faecalis* biofilm matrix is mostly composed of eDNA, polysaccharides, lipoteichoic acid and extracellular proteases (34), but the underlying mechanisms leading to its formation remain to be clearly established.

Recent studies revealed a vast remodelling of metabolic pathways during the early phase of biofilm formation in *Bacillus subtilis* and *B. cereus,* including modulation in fermentation processes, energy production, primary and secondary metabolism pathways (e.g. fatty acid, carbon, amino acid and nucleotides metabolism). It is only recently that these pathways have been involved in biofilm formation (35, 36). In *S. aureus* or *E. faecalis*, it has been reported that metabolic pathways such as the Krebs cycle (37), secondary metabolites production (38) and respiration (39, 40) are positively or negatively regulated during biofilm formation. Identifying and understanding the cellular adaptation that bacteria needs to perform when forming a biofilm could lead to the identification of new therapeutic targets to inhibit this process. Such targets would be of great help in designing therapeutic strategies to treat various chronic infections, to improve the effectiveness of current drugs and limit the development of resistance.

Here, we used a transcriptomic approach to determine if certain cellular pathways were similarly regulated during biofilm formation in *S. aureus* and *E. faecalis.* Our analysis showed that, depending on the strain examined, between 242 and 466 genes were differentially expressed (DEGs) between planktonic and biofilm conditions. Our gene set enrichment analysis (GSEA) revealed that biosynthesis of secondary metabolites, biosynthesis of antibiotics and purine biosynthesis are three commonly upregulated metabolic pathways in biofilm formation. Of note, there were no pathway similarly downregulated between *S. aureus* and *E. faecalis* during biofilm formation. Purine biosynthesis and in particular the *de novo* IMP biosynthesis pathway were the most enriched cellular function that arose from our analysis. Gene disruption and drug targeting in the *de novo* IMP biosynthesis pathway revealed its essentiality for robust biofilm growth.

## Results

### Determination of the growth conditions for biofilm harvest

Biofilms formed by various strains of *S. aureus* and *E. faecalis* can differ in their compositions in a natural environment or *in vitro* (34, 38). Consequently, to ensure that our assay includes commonly regulated cellular pathways in a strain-independent manner, we analysed two strains of each pathogen. For *S. aureus* we used the MRSA strain USA300 and the MSSA strain SH1000, which produce PIA-independent and PIA-dependent biofilm respectively (22, 41). Since differences in biofilm formation between *E. faecalis* strains are not known, we used *E. faecalis* V583 and ATCC 29212, two widely used reference strains. A core genome analysis of these two *E. faecalis* strains using PATRIC Proteome Comparison Service (42) revealed that they share an average of 95.4% of protein sequence identity between their coding DNA sequence (CDS) and 1082 conserved genes with 100% of sequence identity. *S. aureus* strains share an average of 98.8% sequence identity and 2250 conserved genes with 100% of sequence identity (see Fig. S1A and Table S5). Since the two *S. aureus* strains produce very distinct biofilms while sharing a greater identity between their genes than the two *E. faecalis* strains, we expected to find differences in biofilm formation between the *E. faecalis* strains. Interspecies comparisons showed that the two *S. aureus* and *E. faecalis* strains have an average of 43.3% of sequence identity between their CDS (see Fig. S1A and Table S5). The 459 genes with at least 50% homology between the four strains of this study are involved in different metabolic pathways, such as amino acid, carbon, fatty acid or nucleotide metabolism (Fig. S1B), suggesting that they could be conserved between the two species.

We aimed at analyzing mRNA during the early stage of biofilm development, which coincides with the most pronounced biofilm cellular adaptations in other Gram-positive species (35, 36). However, since this phase might be reached at different time by the different strains, we monitored biofilm formation using crystal violet to determine the best time point to harvest the mRNA. As biofilms of *S. aureus* and *E. faecalis* reached their maximum development after 9h and 7h respectively (Fig 1), cells from both strains were harvested just before this plateau. In parallel, we monitored planktonic cell growth (Optic Density at 600nm; OD_600_) in the same conditions but with agitation to harvest the cells just before they reached their growth plateau. These procedures ensured that planktonic and biofilm cells were at similar growth stage when compared. Of note, we also used two different media, BHIg(s) and TSBg(s), known to induce *S. aureus* and *E. faecalis* biofilm formation and to potentially influence biofilm matrix composition (44–46). Growth dynamic in BHIg and TSBg were comparable, except for *S. aureus* USA300, which formed more abundant biofilm in BHIg.

**Figure 1.**
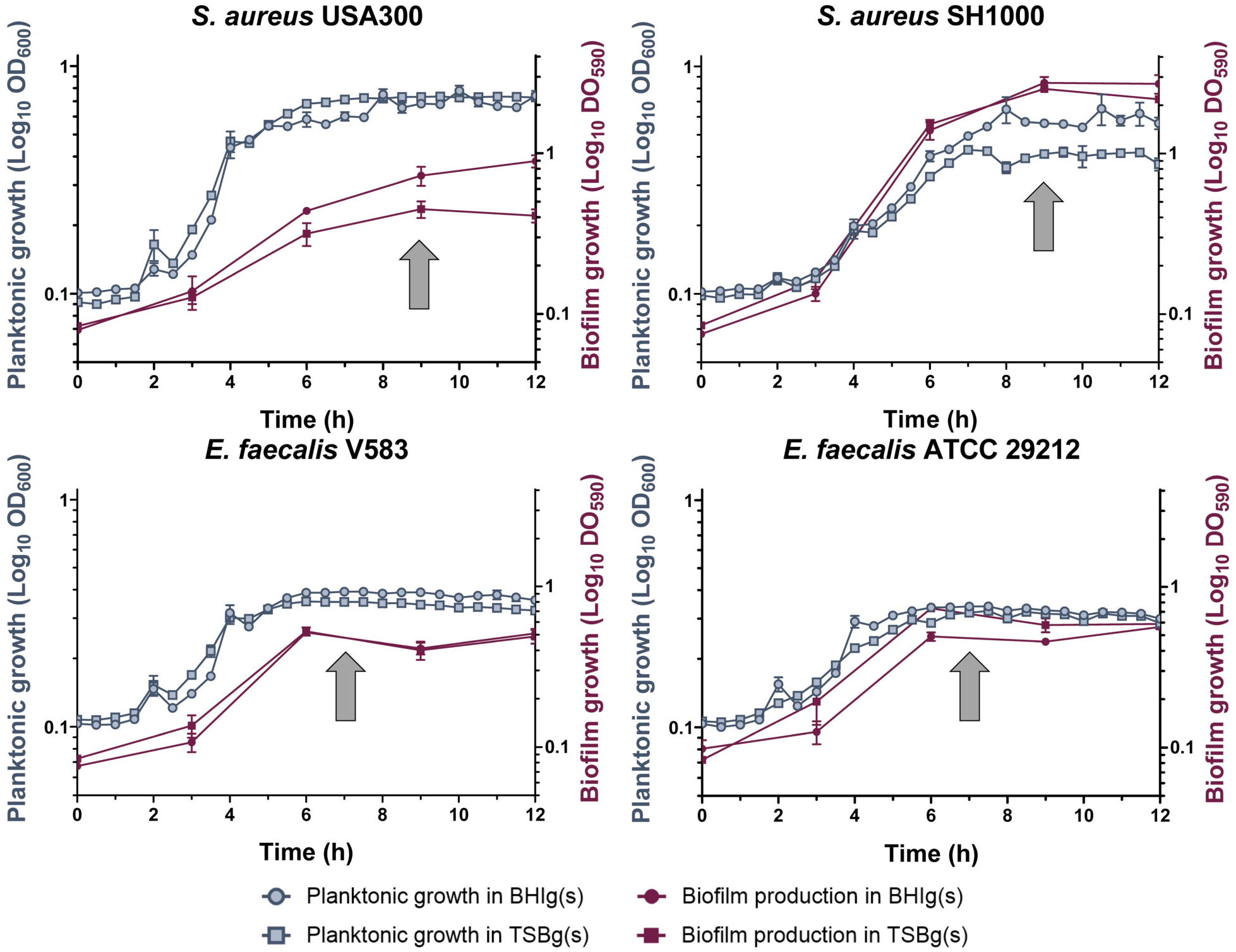
Growth dynamic of *S. aureus* and *E. faecalis* strains in a planktonic state or during biofilm formation. Planktonic growth (in blue) was performed at 37°C in shaking condition and measured by OD_600_, while biofilm growth (in purple) was performed at 37°C without shaking and measured by crystal violet assay (OD_590_). Growth was observed in either BHIg(s) (circle) or TSBg(s) (square). Each point represents the mean of at least 3 biological replicates ± standard deviation (SD). RNA harvesting time is indicated by a gray arrow.

### Transcriptomic analysis reveals global changes in gene expression during biofilm formation

To determine changes in gene expression levels, we used an RNA-seq approach on cells harvested from planktonic and biofilm conditions from biological triplicates of each experimental condition that were pooled together. A differential expression analysis of principal component analysis (PCA), showed that they mostly clustered together (see Fig. S2). Genes were considered differentially expressed in biofilm versus planktonic when showing a statistical significance (adjusted *P-*value < 0.05) and a fold change (log2 ratio) ≥ |2.0|. All media combined, 466 unique differentially expressed genes (DEGs; ~17.3% of the genome) were found for *S. aureus* USA300, 280 unique DEGs (~9.6% of the genome) for *S. aureus* SH1000, 242 unique DEGs (~7.7% of the genome) for *E. faecalis* V583 and 460 unique DEGs (~16.4% of the genome) for *E. faecalis* ATCC 29212 (data are available in the Gene Expression Omnibus (GEO) database under the accession number GSE162709). Upregulated (log2 fold change ≥ 2.0) and downregulated (log2 fold change ≤ −2.0) DEGs distribution among *S. aureus* and *E. faecalis* strains and media are displayed in the Volcano plots shown in Fig. 2.

**Figure 2.**
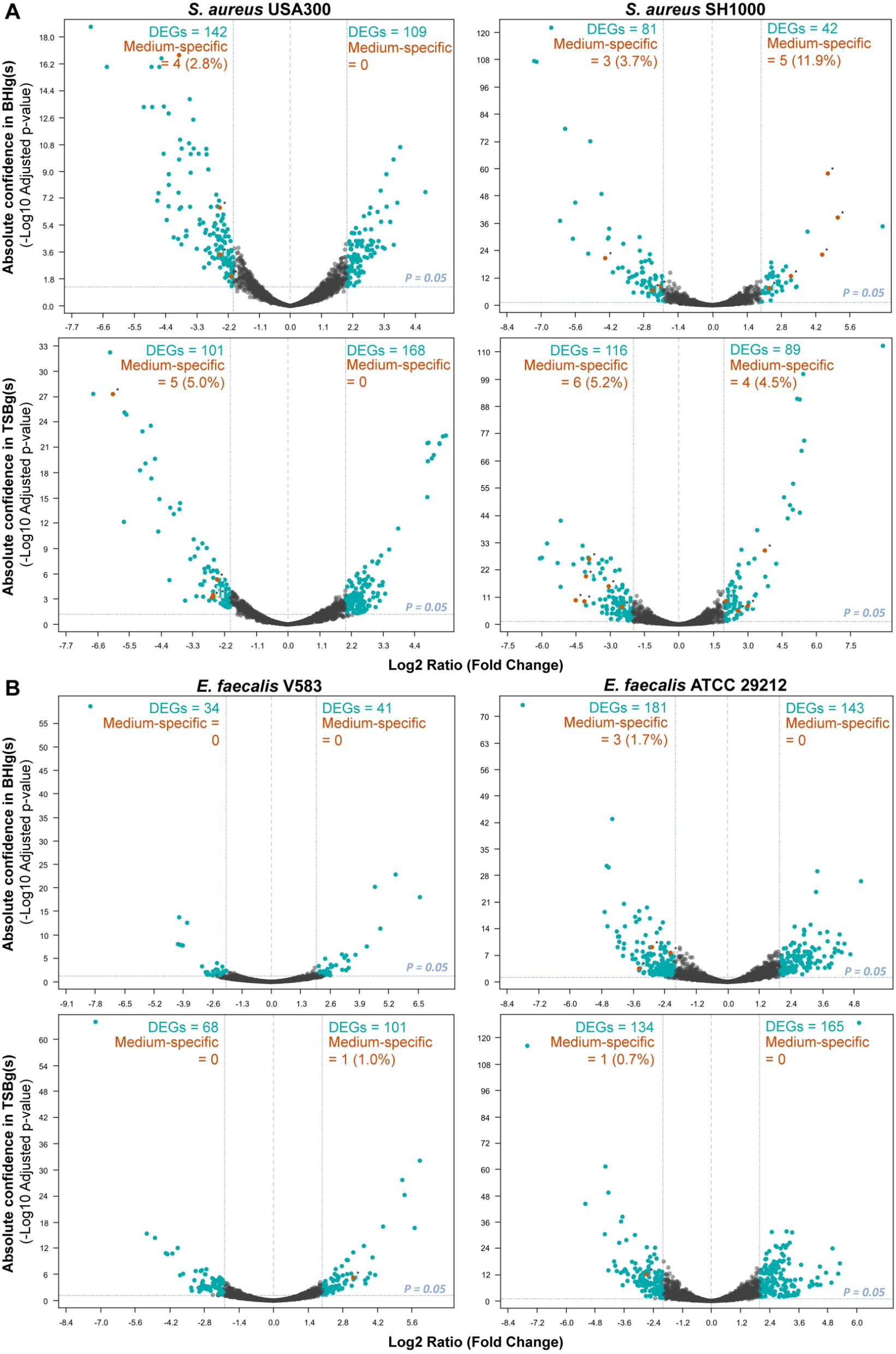
Transcriptional profiles of biofilm-grown cells and planktonic cells. The DEGs (blue dots) adjusted *P*-value cut-off was 0.05. Genes were considered up-regulated in biofilm if the log2 ratio (fold change) was ≥ 2.0 and down-regulated in biofilm if the fold change was ≤ −2.0. DEGs from the media effect transcriptome analysis were then compared to the DEGs from the biofilm growth transcriptome analysis. Genes found in both analyses are represented in orange dots and represent possible DEGs influenced by media instead of biofilm formation. (A) *S. aureus* USA300 and *S. aureus* SH1000 DEGs in TSBg(s) or BHIg(s). (B) *E. faecalis* V583 and *S. aureus* 29212 DEGs in TSBg or BHIg.

### Media effect on gene expression

Metabolic pathways have been previously shown to be involved in biofilm formation in several Gram-positive bacteria (35, 36, 43), but these pathways can also be modulated by nutrients in the growth medium. Thus, we examined the influence of two different biofilm growth media on gene expression patterns to determine if the regulation we observed was specific to the growth medium used or to biofilm formation. To do that, we determined genes that were differentially expressed in TSBg compared to BHIg, all growth condition combined. For *S. aureus,* 75 genes of USA300 and 149 genes of SH1000 had their expression significative modulated by the two media. Of these, a total of 9 and 18 were also found differentially expressed in biofilms for USA300 and SH1000, respectively (Fig. 2). For *E. faecalis* strains, we observed that only 11 genes of V583 and 31 genes of ATCC 29212 were differentially expressed in one or the other media, all growth conditions combined. Consequently, the growth medium used had little to no effect on the gene expression pattern of *E. faecalis* strains during biofilm formation, with less than 1% of all DEGs being also differentially expressed when comparing the transcriptomes obtained from different growth media. Altogether, the biofilm-inducing medium used had only a minor influence on the pattern of gene expression during biofilm formation.

### Gene set enrichment analysis reveals global cellular adaptations during biofilm formation

We compared biofilm and planktonic gene expression patterns from the four strains in TSBg(s) or BHIg(s) using DAVID Bioinformatic resources (47). Since *S. aureus* USA300 and *E. faecalis* ATCC 29212 locus tags were not recognized by DAVID, we attributed the corresponding recognized locus tag of their species counterpart based on sequence homology and orthologs with the help of BioCyc Pathway tools version 23.0 (48) and AureoWiki (49). Gene up- or downregulated in the heatmap of *S. aureus* and *E. faecalis* represented 497 and 316 GO: BP (Gene Ontology Biological Processes) terms, respectively (Fig. 3A & B). The 20 most up- and down-regulated GO: BP terms for each species, all strains and media combined, are shown in Panel C of Fig. 3. The same analysis performed with the Kyoto Encyclopedia of Genes and Genomes (KEGG) pathways as the annotation source showed that 103 KEGG functions of *S. aureus* and 97 KEGG functions of *E. faecalis* were represented in up- or down-regulated genes (Fig. S3). Many metabolism-related pathways were up-regulated during biofilm formation, all annotation sources combined, such as carbon, fatty acid, nucleotides and some amino-acid metabolism genes.

**Figure 3.**
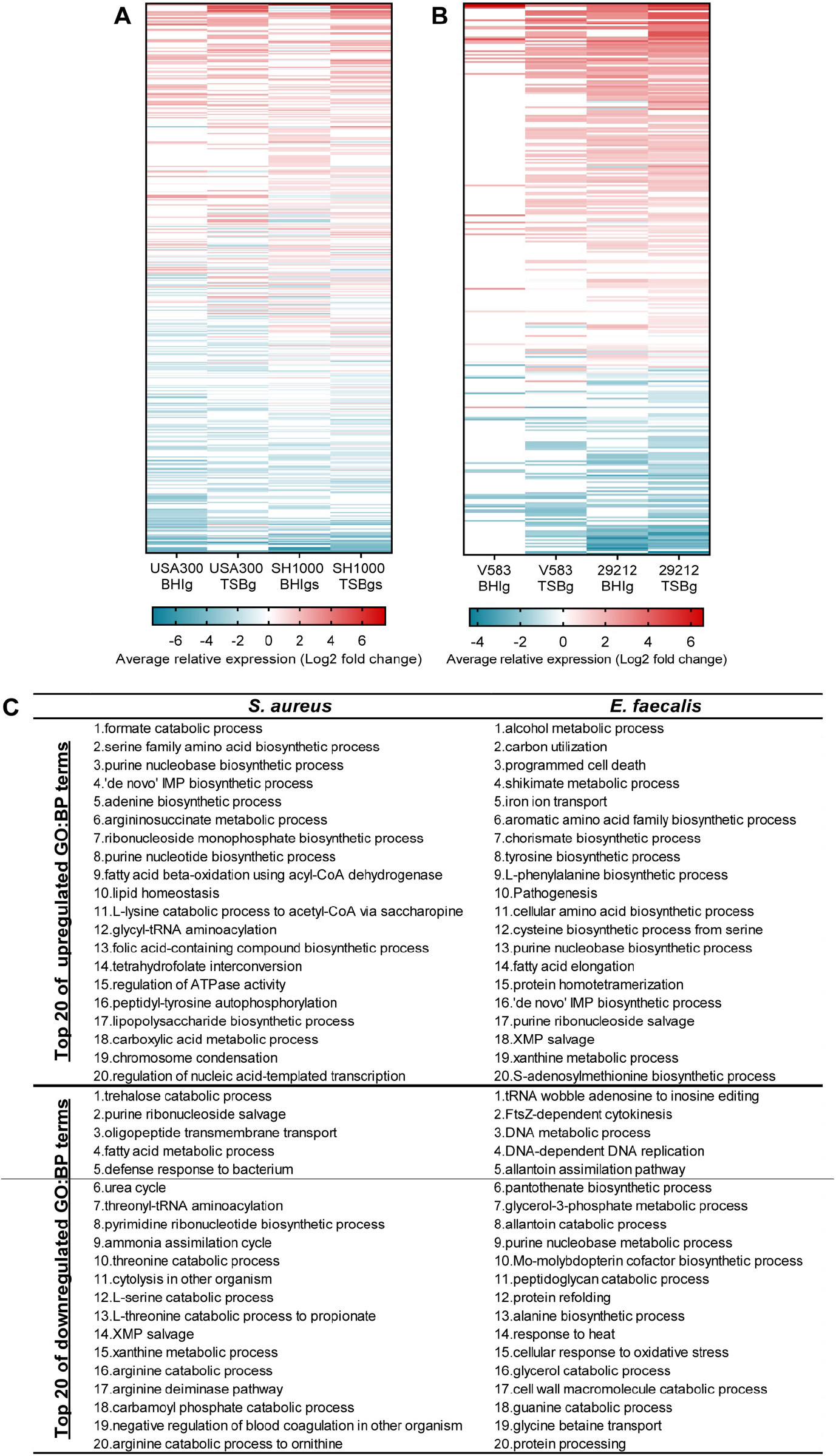
Average relative expression of GO:BP terms during biofilm formation. Each row of both heatmaps represents a single functional annotation term from DAVID Bioinformatic software using GO:BP annotation source. A term’s relative expression is the average of the log2 fold change of every associated gene with an adjusted *P*-value ≤ 0.05 from the transcriptomic analysis. The red (up-regulated) and blue (down-regulated) colour scale indicates relative expression level in average log2 fold change. (A) Functional annotation terms’ (total = 497) relative expression in *S. aureus* USA300 and SH1000, in BHIg(s) or TSBg(s) medium. (B) Functional annotation terms’ (total = 316) relative expression in *E. faecalis* V583 and 29212, in BHIg or TSBg medium. (C) Table of the top 20 up-regulated and down-regulated GO:BP terms for each species, by calculating the average of the relative expression of each term for the 2 strains in each medium.

### Common cellular adaptations between S. aureus and E. faecalis strains during biofilm formation

Next we determined among the cellular functions that were up- and downregulated during biofilm formation, which ones were common to all strains and media used. We attributed annotation with a GSEA using the DAVID Bioinformatics database and GO: BP, KEGG pathways, Gene Ontology Molecular Function (GO:MF) and UniProt keywords (Up_Kw) were used as annotation sources to increase the range of finding potential functions. As shown in Fig. 4A, only eight terms from all annotation sources were upregulated in all strains and growth media during biofilm formation. Interestingly, the 3 most enriched terms were related to purine biosynthesis. This result points toward a conserved role of the purines biosynthesis pathway during biofilm formation in *S. aureus* and *E. faecalis*. Other cellular pathways identified as commonly upregulated in biofilms were the biosynthesis of antibiotics and the biosynthesis of secondary metabolites. However, the core genome analysis of the four strains showed that specific genes involved in these pathways were not conserved between *S. aureus* and *E. faecalis* (Fig. S1B). Our analysis did not highlight commonly downregulated terms during biofilm formation. As depicted in Fig. 4B, although purines biosynthesis was upregulated in biofilm formation between *S. aureus* strains, pyrimidine biosynthesis and *de novo* UMP biosynthetic process were downregulated. Between *E. faecalis* strains, only the phosphotransferase system (PTS) was identified as being commonly downregulated in biofilm formation. To eliminate bias or omission that could be due to usage of a single GSEA tool, we submitted the same DEGs to ShinyGO v0.61 (50) and confirmed that purine biosynthesis was strongly regulated during biofilm formation (Fig. S4).

**Figure 4.**
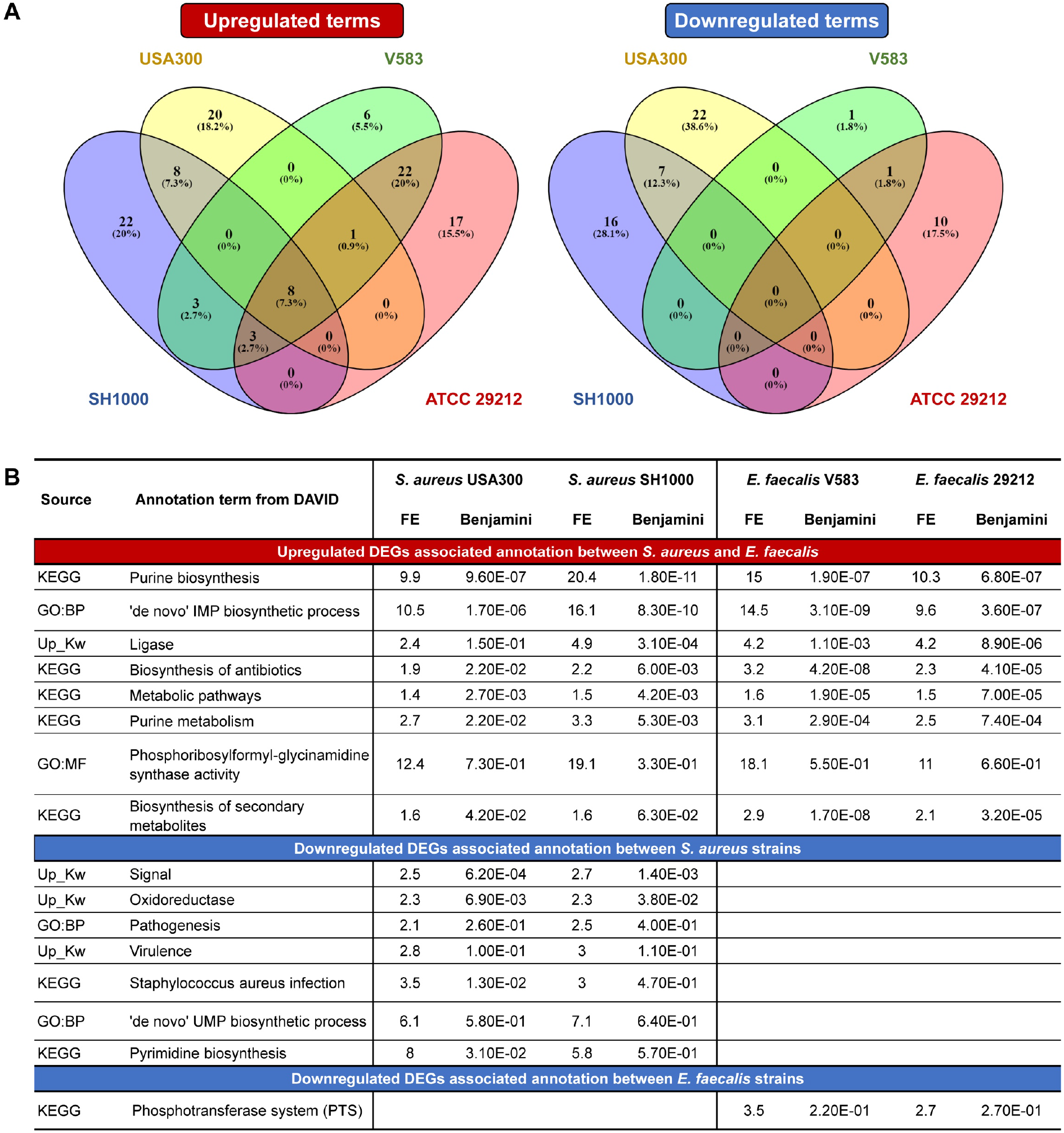
Common cellular adaptation between *S. aureus* and *E. faecalis* strains. (A) Venn diagram representation of the unique and common functional terms of the upregulated DEGs (*P*-value ≤ 0.05, log2 fold change ≥ 2.0) and downregulated DEGs (*P*-value ≤ 0.05, log2 fold change ≤ −2.0) between *S. aureus* USA300 (yellow), *S. aureus* SH1000 (blue), *E. faecalis* V583 (green) and *E. faecalis* 29212 (red). (B) Table of the common cellular pathways between the four strains of this study, either upregulated or downregulated. GSEA was performed with DAVID Bioinformatic resources using KEGG pathways, GO:BP, GO:MF and Up_keywords annotation source. Pathways with a fold enrichment (FE) superior to 1.5 are considered interesting pathways while confidence in the enrichment is indicated by the Benjamini value.

### Purine biosynthesis remodelling during biofilm formation

Bacteria can metabolize purines *de novo* from 5’-phosphoribosylamine-1-pyrophosphate (PRPP) through biosynthesis of inosine monophosphate (IMP), or they can recycle them from the nucleic acids present in their environment via the purine salvage pathway (51). These metabolic pathways, with each gene involved in these reactions next to the arrow, are depicted In Fig. 5. In adjacent boxes we report their expression level in biofilm (log2 fold change), for all media combined. Overall, genes in the *de novo* pathway were homogeneously strongly upregulated (log2 fold change ≥ 2.0) in all *S. aureus* and *E. faecalis* strains. Only *purB,* the sole gene found also in the salvage pathway, did not show a strong up-regulation. On the other hand, gene expression levels of the biosynthesis and salvage pathways were heterogeneously and weakly regulated.

**Figure 5.**
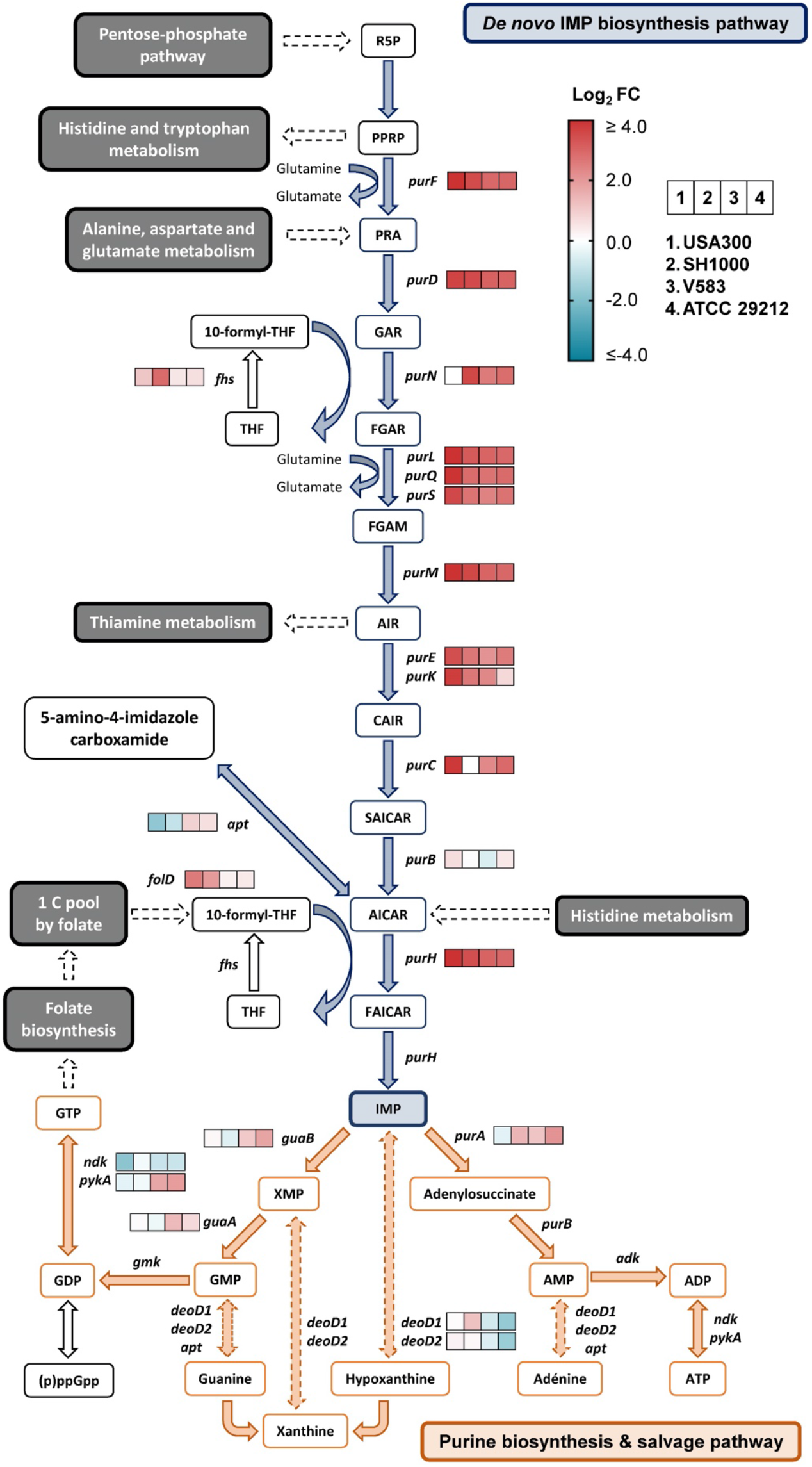
The *de novo* IMP biosynthesis pathway is strongly upregulated during biofilm formation. The purine biosynthesis pathway includes the *de novo* IMP biosynthesis pathway (in blue) and the purine salvage pathway (in orange). Plain arrows indicate a unique reaction while dotted arrows indicate multiple reactions. The expression level (log2 fold change) during biofilm formation of the genes involved in the pathway (all media combined), as obtained by the transcriptomic analysis, are indicated by a 4-square stripe (from left to right: *S. aureus* USA300, *S. aureus* SH1000, *E. faecalis* V583 and *E. faecalis* 29212). The red and blue colour scale indicates high and low expression level, respectively. For abbreviations, see Table S3.

### Genes involved in the de novo IMP biosynthesis pathway are important for biofilm production

Since IMP biosynthesis was found highly up-regulated in biofilm for all the strains of this study, we wanted to confirm that this pathway is important for biofilm formation. Thus, we performed biofilm growth assays using *S. aureus* USA300 strains with transposon insertion in *purF, purD, purL, purQ, purS, purM, purK* and *purH* genes from the *de novo* biosynthesis pathway (Nebraska Transposon Mutant Library; NTML) (52). We also used a strain with a transposon in *purA* from the purine biosynthesis and salvage pathway, in *purB* from both pathways and in *folD* from the 1 C pool by folate pathway. The bifunctional protein FolD synthetizes 10-formyl tetrahydrofolate (10-formyl-THF), a cofactor used by *purN* and *purH* during *de novo* IMP biosynthesis (53). Finally, we used strains deleted for *deoD-1* and *deoD-2* from the purine salvage pathway. As shown in Fig. 6A, disruption of genes involved in the *de novo* pathway significantly impaired biofilm production, while interrupting genes of the biosynthesis and salvage pathway did not affect (*purA, deoD-1, deoD-2*) or only slightly impaired (*purB*) biofilm production. Inactivation of genes involved in the latest step of IMP showed more defects in biofilm production, with *purN, purL, purQ, purH* and *purM* having the most significant impact. Importantly, the effects observed on biofilm production by purine biosynthesis mutants were not due to a delayed or decreased cell division rate (Fig. 6B). Only *purA* and *purB* mutants showed a significant decrease in planktonic growth during the late exponential phase of growth (~ 9h), which could partly explain their mild effect on biofilm production. However, USA300 biofilm formation relies heavily on eDNA, thus possibly exacerbating the necessity of a functional purine biosynthesis pathway. To examine if the importance of this pathway was conserved in PIA-dependent biofilms, we treated SH1000 with ATIC dimerization inhibitor, a drug targeting the bi-functional enzyme PurHJ. As shown in Fig. 6C, this drug successfully reduced biofilm formation by SH1000 without affecting cell growth. While the high concentration of drugs required is likely due to the fact that the ATIC inhibitor was developed specifically for cancer treatment in mammalian cells, this result supports that *de novo* purine biosynthesis pathway constitutes an interesting target to reduce biofilm formation.

**Figure 6.**
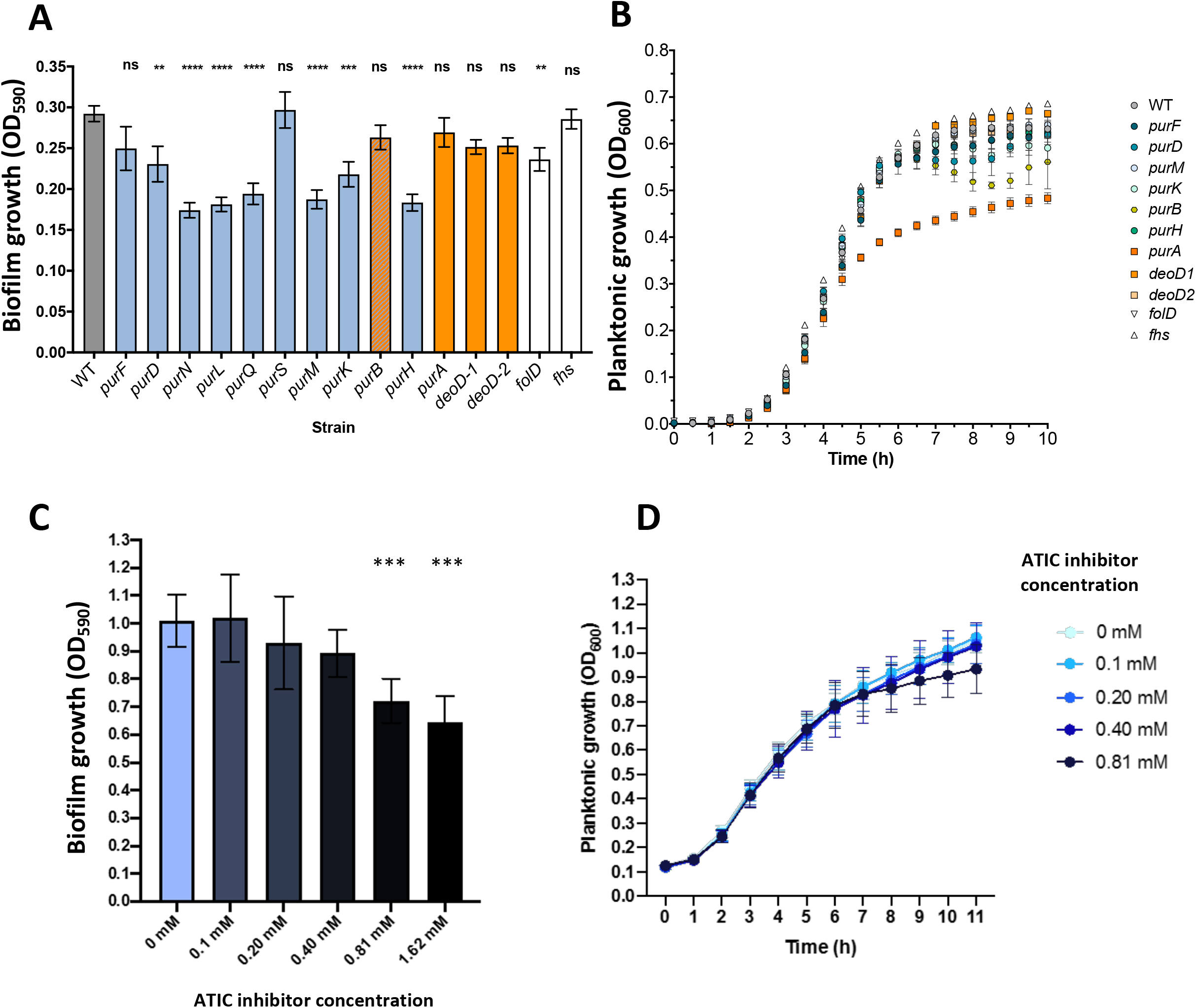
Inhibition of *de novo* purine biosynthesis pathway affect biofilm formation. (A) Biofilms of *S. aureus* USA300 containing transposon insertion (NTML) in gene encoding for the purine biosynthesis pathway were formed in TSBg and quantified by crystal violet. Each bar represents biofilm production (in OD_590_) of at least 3 biological replicates ± SEM (one-way ANOVA; ns, not significant; * *P*-value < 0.05; ** *P*-value < 0.01; **** *P*-value < 0.0001). (B) Planktonic growth of *S. aureus* USA300 mutants in TSBg. Each point of the curve represents the cell density (OD_600_) of at least 3 biological replicates ± SEM. (C) Biofilms of *S. aureus* SH1000 were formed in TSBgs containing different concentration of ATIC dimerization inhibitor, and quantified by crystal violet. Each bar represents biofilm production (in OD_590_) of at least 3 biological replicates ± SD (one-way ANOVA; *** *P*-value < 0.001). (D) Planktonic growth of *S. aureus* SH1000 in presence of various concentration of ATIC inhibitor in TSBgs. Each point of the curve represents the cell density (OD_600_) of at least 3 biological replicates ± SD.

## Discussion

Regulatory mechanisms leading to biofilm formation are often species or even strain-specific, making it difficult to develop anti-biofilm strategies that target a wide spectrum of bacteria (59, 60). Recent developments in biofilm research highlighted metabolic remodelling during biofilm formation in some Gram-positive bacteria (35, 36). Using a transcriptomic approach with Gram-positive strains of clinical importance, we reported here a conserved regulation of gene expression in specific metabolic pathways during biofilm formation (Fig. 3).

TSB and BHI medium supplemented with glucose and/or sodium chloride are two frequently used media to induce biofilm formation of *S. aureus* and *E. faecalis* (45, 61). However, it was suggested that TSB promotes PIA production for *S. aureus* while BHI promotes a protein-based matrix (44). According to our transcriptomic analysis, several genes were found differentially expressed in one or the other medium, all strains and growth conditions combined. Many of those DEGs were attributed to primary metabolism pathways and some to secondary metabolism pathways (Table S4). We observed that the growth media used for biofilm formation had some effect on the transcriptome profiles in *S. aureus*, possibly due to environmental factors, but that these profiles were mostly unchanged in *E. faecalis* strains (Fig. 2). Indeed, different growth media were reported to influence biofilm phenotypic properties and matrix composition in other Gram-positive strains such as *B. subtilis* (62). Although the growth medium mildly modulated gene expression during planktonic and/or biofilm growth of some *S. aureus* strains, it did not prevent us from identifying a large number of up-regulated genes belonging to metabolic pathways that are potential targets for new antimicrobial agents. However, when doing expression analysis of biofilm condition has to be chosen carefully in order to avoid identifying genes whose expression is modulated only in a specific medium.

When comparing the functional enrichment of the DEGs during biofilm formation of *S. aureus* USA300 and SH1000 to *E. faecalis* V583 and ATCC 29212, only three biological functions were found commonly up-regulated during biofilm growth: biosynthesis of secondary metabolites, biosynthesis of antibiotics and purine biosynthesis (Fig. 4). Some of the genes and biological functions we found up-regulated in *S. aureus* during biofilm growth (Fig. 3), such as fermentation, nucleotide metabolism and the TCA cycle, were already observed in other transcriptomic, metabolomic or proteomic studies using other *S. aureus* strains (63–67). In *E. faecalis*, only a few metabolic adaptations during biofilm formation have been reported (68, 69). Therefore, our data provide new information formation of *E. faecalis* that could be exploited in further studies and consolidate previous observations on *S. aureus* biofilm formation.

Several studies previously reported purine biosynthesis as being up-regulated in both Gram-positive and Gram-negative bacteria during biofilm formation (35, 43, 70–74). Similar to those studies, we found a strong up-regulation of the *de novo* IMP biosynthetic process during biofilm formation in *E. faecalis* and *S. aureus* strains (Fig. 5). Gene disruption in this pathway, but not in the purine salvage and biosynthesis pathway, led to a decrease in *S. aureus* USA300 biofilm production (Fig. 6). While it is not the strongest upregulated pathway in our study, *de novo* purine biosynthesis is the only pathway upregulated in all our strains and conditions, which justify further investigation. Indeed, the biological importance of *de novo* purine or IMP formation in biofilm warrants further investigations.

Purines are required for eDNA production, an important constituent of the extracellular matrix of *S. aureus* (75), *E. faecalis* (76) and other Gram-positive bacteria such as *B. subtilis* or *B. cereus* (77). However, a study demonstrated that in *B. cereus*, *purD* and *purH* deletions did not affect significantly the level of eDNA in *B. cereus* matrix compared to the wild-type strain (43). Guanosine monophosphate (GMP) and adenosine monophosphate can lead to the production of cyclic-di-GMP and cyclic-di-adenosine monophosphate (AMP) respectively, two molecules that are involved in biofilm formation in several Gram-positive bacteria (78). GMP can also form guanosine pentophosphate (ppGpp) from GDP, which plays a role in biofilm formation of *E. faecalis* (79). Limiting nucleotides or IMP precursors in a biofilm-inducing medium could help better understand purine implication in the biofilm composition. Drugs targeting different steps of the purine biosynthesis pathway could also be used to investigate this phenomenon in other Gram-positive strains that lack efficient transformation tools or an available mutant library.

Altogether, we have shown that the remodelling of specific core metabolism pathways could be shared among many Gram-positive bacteria during biofilm formation. Our results suggest that purine biosynthesis could be a potent anti-biofilm target in a wide spectrum of Gram-positive bacteria.

## Materials and Methods

### Strains and media

Strains used for the transcriptomic analysis are *S. aureus* USA300, *S. aureus* SH1000, *E. faecalis* V583 and *E. faecalis* ATCC 29212 (a kind gift from F. Malouin, Université de Sherbrooke). Biofilm phenotypic assays were performed with *S. aureus* USA300 mutants from the Nebraska Transposon Mutant Library (NTML) (52) (see Table S1 in the supplementary material). Strains were maintained on Tryptic Soy agar (TSA) or Brain Heart Infusion (BHI) agar plates, and for the NTML mutants, 5 μg/ml erythromycin was added to the media. To induce biofilm formation, Tryptic Soy broth or BHI broth supplemented with 0.5% (wt/vol) glucose (TSBg or BHIg) was used for *E. faecalis* strains and *S. aureus* USA300, and TSBg or BHIg supplemented with 3.0% (wt/vol) sodium chloride (TSBgs or BHIgs) was used for *S. aureus* SH1000.

### Growth measurement (planktonic cells and biofilm)

Colonies from overnight cultures on TSA or BHI agar plates at 37°C were suspended in 1 ml of the appropriate biofilm inducing medium. Then, 200 μl of biofilm inducing medium was added in wells of a 48-wells culture plate and inoculated to an OD_600_ of 0.005 with the bacteria suspension. For planktonic growth measurement (at least 3 biological replicates), plates were incubated in a Synergy™ HT plate reader (BioTek, Winooski, VT, USA) at 37°C with fast and continuous agitation to prevent biofilm formation and monitored at 600 nm every 30 min for at least 18h. For Figure 6C and D, ATIC dimerization inhibitor (Millipore-Sigma) was added from a 1 mg/mL stock in distilled water.

For biofilm formation, plates were incubated at 37°C without agitation for 20h; quantification (at least 3 biological replicates) was performed using the crystal violet method (54). Briefly, biofilms were gently washed with phosphate-buffered saline (PBS), covered with 200 μl of 0.01% (wt/vol) crystal violet and incubated for 20 min at room temperature, protected from light. Each well was then gently washed with sterile distilled water to remove the excess of dye. Stained biofilms are solubilized in 200 μl of acetic acid 33% and quantified by spectrophotometry at 590 nm.

### mRNA extraction and isolation

Planktonic cells (3 biological replicates) were washed with cold PBS and kept in RNA stabilization solution (25 mM sodium citrate, 10 mM EDTA, 47% ammonium sulphate, adjusted to pH 5.2) until all samples were harvested. Biofilms (3 biological replicates) were washed with cold PBS and then suspended in RNA stabilization solution by scraping the bottom of the well until enough cells were harvested. Planktonic and biofilms cells were then centrifuged, the supernatant discarded, and the pellet kept at −80°C until RNA extraction.

Cells were suspended in 250 μl of fresh lysis buffer (45 mg/ml of lysozyme, 16 μl/ml of lysostaphin 5 mg/ml and 10 μl/ml of mutanolysin 5 U/μl in Tris-EDTA) and incubated for 1h at 37°C, while mixed by every 15 min. They were then transferred in O-Ring tubes with 25 to 50 mg of acid-washed beads and mixed with 3 volumes of TRI Reagent®, to be lysed mechanically with a FastPrep-24™ Classic (MP Biomedicals, Santa Anna, CA, USA) (speed of 4.5 m/sec, 2x 20s). Total RNA was extracted with the Direct-zol™ RNA Miniprep Plus kit (Zymo Research, Irvine, CA, USA) following the manufacturer’s instruction, including the DNAse I (New England Biolabs, Ipswich, MA, USA) treatment in the extraction column. RNA quantity and the integrity number (RIN) were evaluated with an Agilent 2100 Bioanalyzer (Agilent Technologies, Santa Clara, CA, USA). To remove ribosomal RNA (rRNA), samples were treated using a Pan-Prokaryote riboPOOL™ kit (siTOOLs Biotech Gmbh, Planegg, Germany) coupled with streptavidin magnetic beads (New England Biolabs, Ipswich, MA, USA). mRNA was eluted in 50 μl of nuclease-free water and then cleaned and concentrated using a Clean & Concentrator kit (Zymo Research, Irvine, CA, USA). mRNA samples were kept at −20°C if used on the same day or frozen at −80°C for long-term storage. Ribo-depletion success was assessed by verifying the diminution or disappearance of the 16s rRNA and 23s rRNA peaks with an Agilent 2100 Bioanalyzer, between a treated sample and a non-treated sample.

### Construction of prokaryotic Illumina libraries

Libraries were prepared for RNA sequencing according to the instructions of the NEBNextl® Ultra™ II Directional RNA Library Prep Kit for Illumina® (New Englands Biolabs, Ipswich, MA, USA), with the following exception. RNA samples were fragmented 7 min instead of 15 min to produce fragments longer than 100 pb. Instead of the adaptors provided with the kit, we used TruSeq compatible YIGA adaptors (see Table S2 in the supplementary material). Nucleic acid purification was performed using Mag-Bind® TotalPure NGS beads (Omega Bio-Tek Inc, Norcross, GA, USA) following the recommended ratio of the NEBNext® kit. Ligation products were subject to PCR amplification using a reverse primer containing a unique 8 bp index for each sample (see Table S2 in the supplementary material). The concentration of nucleic acid in the libraries was measured using a Quant-iT™ PicoGreen™ dsDNA Assay Kit (Invitrogen, Carlsbad, CA, USA) and 1.13 ng of each sample were pooled in 80 μl of nuclease-free water. Quantity and quality of the final mix were confirmed with an Agilent 2100 Bioanalyzer before submitting to a Nextseq500 High Output 75 bp sequencing.

### RNA sequencing data analysis

Approximately 8,000,000 reads per library were obtained using an Illumina Nextseq system providing single-ended reads of 75 bp each. The data was analyzed using Geneious Prime 2020.0.3 (https://www.geneious.com). Fastq files were trimmed with the BBDuk plugin using the default parameters and eliminating reads shorter than 10 bp. Trimmed data were then aligned with Bowtie2 (55) using an “end-to-end” alignment type, a “High Sensitivity / Medium” preset and using the “do not trim option”, against the following reference chromosomes: GenBank accession number GCA_000013425.1 (for *S. aureus* SH1000, a close descendent of the ancestral strain NCTC 8325 (56)); GenBank accession number GCA_000017085.1 (for *S. aureus* USA300); GenBank accession number GCA_000742975.1 (for *E. faecalis* ATCC 29212) and GenBank accession number GCA_000007785.1 (for *E. faecalis* V583). Finally, the differentially expressed genes (DEGs) were assessed with DESeq2 (57) by combining the 3 biological replicates of each condition and comparing biofilm cultures vs planktonic cultures in either TSBg(s), BHIg(s) or both. Only the genes with an adjusted *P*-value ≤ 0.05 were retained for further analysis. DEGs were scored as up-regulated if they had a log2 fold change ≥ 2.0 and down-regulated if they had a log2 fold change ≤ −2.0. For the transcriptomic study on growth media effect on genes expression, DEGs with a log2 fold change ≥ 2.0 were considered up-regulated in BHIg(s) while DEGs with a log2 fold change ≤ 2.0 were considered up-regulated in TSBg(s).

### Functional annotation

The various gene set enrichment analysis (or functional annotation) were performed using DAVID Bioinformatics Resources version 6.8 (47). The locus tag of the upregulated and downregulated DEGs were submitted as a list to DAVID and then analyzed with the Functional Annotation tool. The UP_KEYWORDS of the Functional_category, the KEGG_PATHWAY of the Pathways category, the GOTERM_BP_DIRECT and/or GOTERM_MF_DIRECT of the Gene_Ontology category were selected as possible annotation sources. The analysis was performed with the functional annotation chart, including the default thresholds of a count of 2 and an EASE of 0.1 and applying the following statistical test: Fold Enrichment (FE), Benjamini and FDR. Following DAVID recommendations, function with a fold enrichment ≥ 1.5 and a benjamini value < 0.05 were considered interesting. The GSEA comparing biofilm vs planktonic conditions was confirmed with the ShinyGO v0.61 software. The locus tags of the same DEGs were submitted as a list, using either the *E. faecalis* STRINGdb or the *S. aureus* NCTC8325 STRINGdb and the default *P*-value cut-off (FDR) of 0.05. Both KEGG pathways and Gene Ontology: biological process (GO:BP) were considered. To assess the common pathway between the 4 strains used in this study, their list of all the enrichment terms from DAVID or ShinyGO was compared by the Venn diagram generator online tool Venny 2.1.0 (58).

### Core genome circular comparison

To identify potential conserved cellular pathways among the strain used in this study, each of them was submitted to the Proteome Comparison Service performs, a protein sequence-based genome comparison tool from PATRIC (42), using the default parameters and the following reference genomes: *Staphylococcus aureus* subsp. *aureus* NCTC 8325 (Genome ID: 93061.5), *Staphylococcus aureus* subsp. *aureus* USA300_TCH1516 (Genome ID: 451516.9), *Enterococcus faecalis* V583 (Genome ID: 226185.9) and *E. faecalis* ATCC 29212 (Genome ID: 1201292.8). Genome comparison was illustrated in a scalable vector graphic (SVG) image file produced by Circos. All the recognized locus tags of the genes sharing over 50% of sequence identity in the genome comparison text file of every strain were subjected to a GSEA as described earlier.

### Statistical analysis

Statistical analysis was performed using GraphPad Prism 8.4.0. Comparisons were done using Student’s t-test or one-way analysis of variance (ANOVA) followed by Tukey’s multiple-comparison test, both with 95% confidence intervals.

### Data availability

All sequence data have been uploaded to the Gene Expression Omnibus under the accession number GSE162709 (https://www.ncbi.nlm.nih.gov/geo/query/acc.cgi?acc=GSE162709)

## Abbreviations

10-formyl-THF: 10-formyl tetrahydrofolate
AMP: adenosine monophosphate
ANOVA: one-way analysis of variance
ALT: antibiotic lock therapy
CDS: coding DNA sequence
DEGs: differentially expressed genes
eDNA: extracellular DNA
EPS: extracellular polymeric substances
FE: Fold Enrichment
GEO: Gene Expression Omnibus
GMP: Guanosine monophosphate
GO:BP: Gene Ontology Biological Processes
GO:MF: Gene Ontology Molecular Function
GSEA: gene set enrichment analysis
IMP: inosine monophosphate
KEGG: Kyoto Encyclopedia of Genes and Genomes
MIC: minimal inhibitory concentration
NIH: National Institutes of Health
NSERC: Natural Sciences and Engineering Council of Canada
NTML: Nebraska Transposon Mutant Library
OD: Optic Density
PBS: phosphate-buffered saline
PCA: principal component analysis
PIA: polysaccharide intercellular adhesin
ppGpp: guanosine pentophosphate
PRPP: 5’-phosphoribosylamine-1-pyrophosphate
PTS: phosphotransferase system
RIN: RNA quantity and the integrity number
rNA: ribosomal RNA
SARM: *S. aureus* resistant to methicillin
SASM: *S. aureus* sensitive to methicillin
SVG: scalable vector graphic
Up_Kw: UniProt keywords.

## Acknowledgments

We thank F. Malouin and J.-P. Côté for their kind gift of strains, L.-C. Fortier and S. Rodrigue for critical reading of the manuscript, and members of the Beauregard, Burrus, Malouin and Rodrigue labs for helpful discussions. This work was supported by a Canada Graduate Scholarships (Master’s Program) from the Natural Sciences and Engineering Council of Canada (NSERC) to M.G. This project was funded by Université de Sherbrooke and FRQ-NT grant 253077.

---

**Figure S1.**
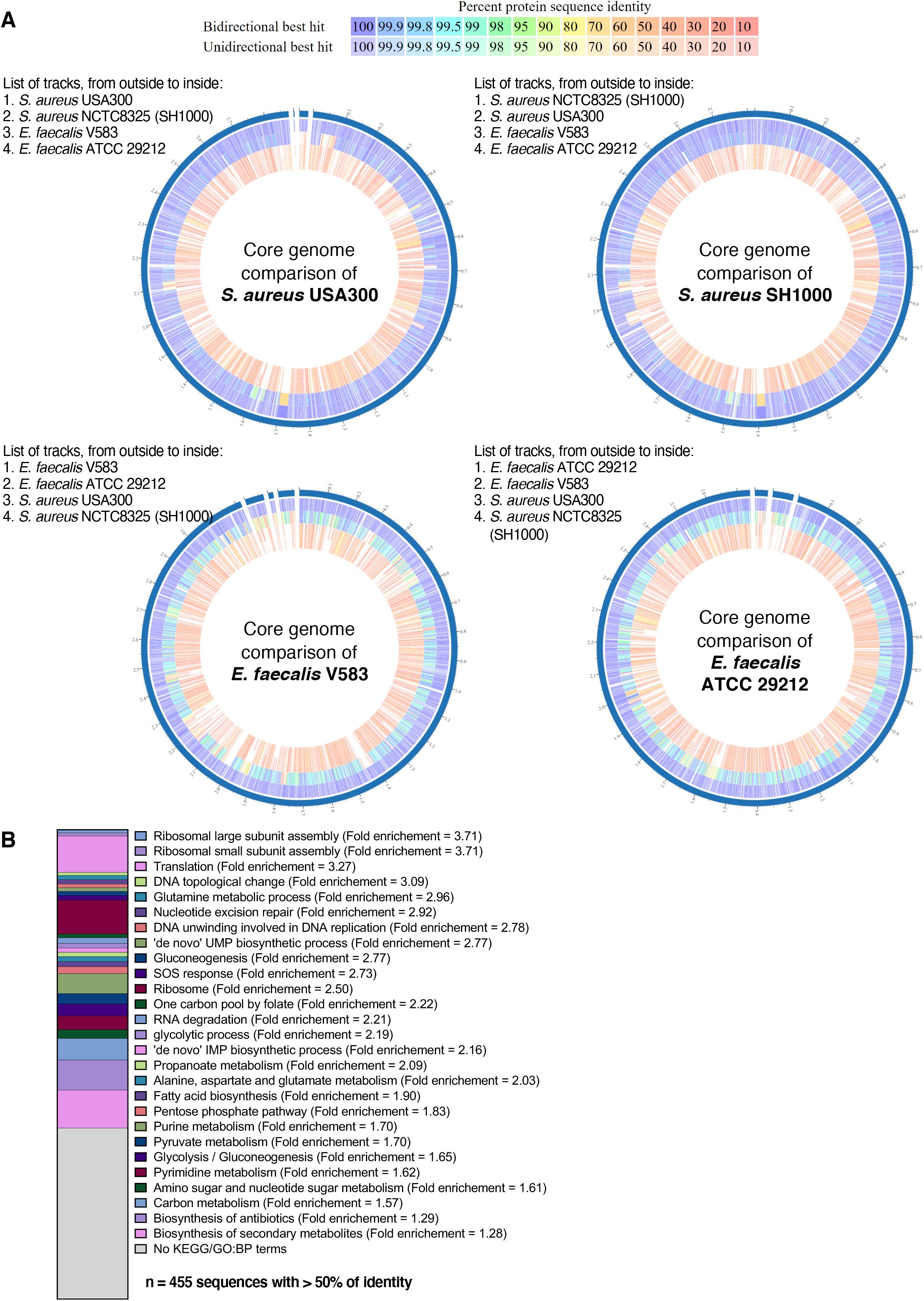
Core genome analysis between Gram-positive strains in a protein sequence-based homology manner. **(A) Circular representation of *S. aureus* and *E. faecalis* strains genomic comparison. Each ring represents the chromosome of a strain, where the outer ring is the reference genome and the three inner rings are the chromosomes of comparison, as indicated on the upper-left corner of each plot. Each line represents a CDS and its % of identity with the reference genome is represented by the blue (high) to the red (low) colour scale. Plots were generated using CIRCOS circular genome data visualization with PATRIC’s proteome comparison service.** (B) Cellular functions from the genes sharing at least 50% of identity (n = 455) in their protein sequence between the 4 strains of the transcriptomic study, obtained with **PATRIC’s proteome comparison service**. Each row from the graph represents the proportion of genes attributed to a function relative to the total number of genes. The fold enrichment of each term is indicated next to its name. Functional annotation was performed with DAVID Bioinformatic software using GO:BP and KEGG pathways as annotation sources.

**Figure S2.**
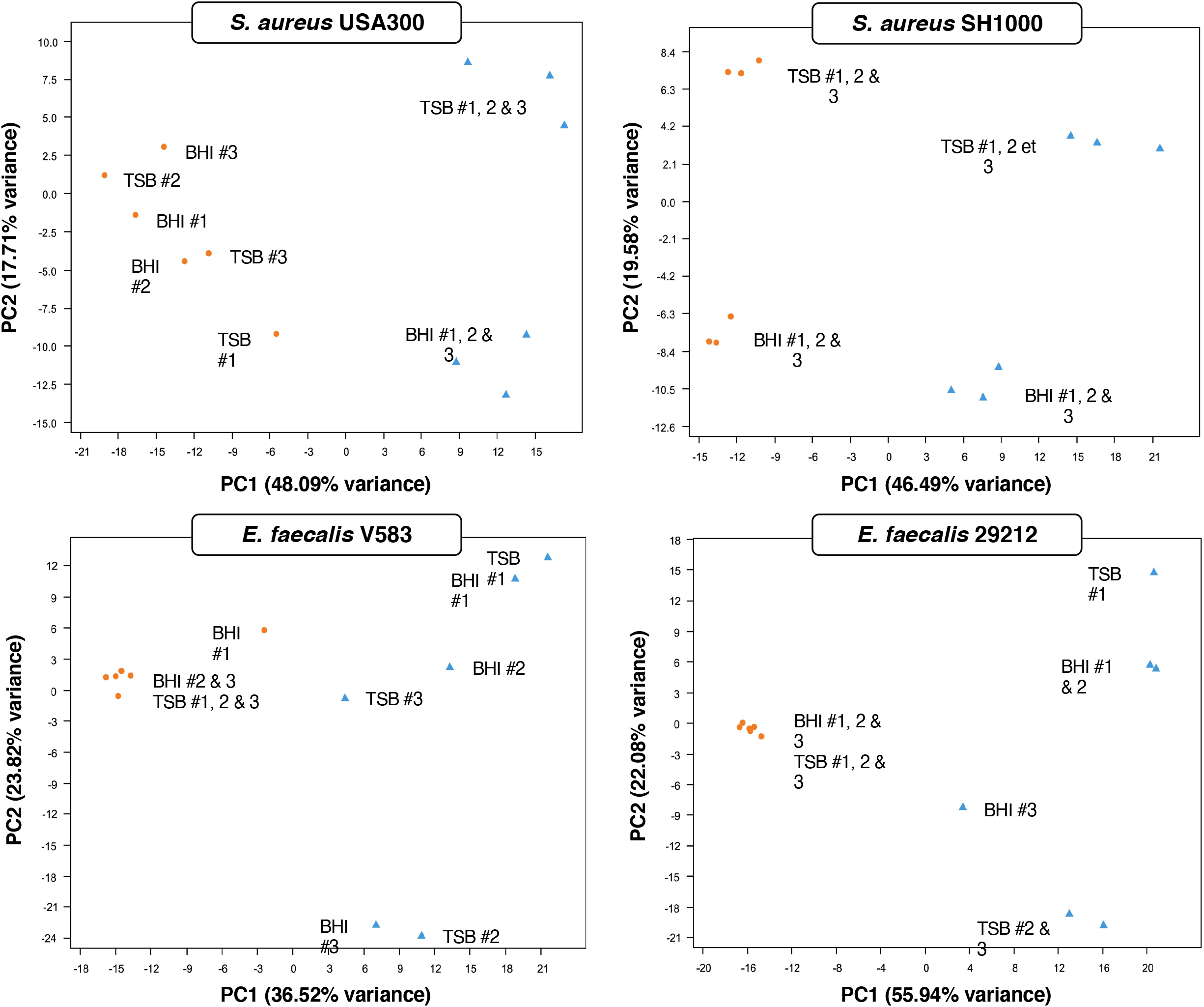
Principal component analysis (PCA) of the variance between the biological replicates from the transcriptomic analysis. The blue triangles represent planktonic cultures while the orange circles represent biofilm cultures of *S. aureus* or *E. faecalis*. The medium and the number of the biological replicate are indicated next to the geometrical icon.

**Figure S3.**
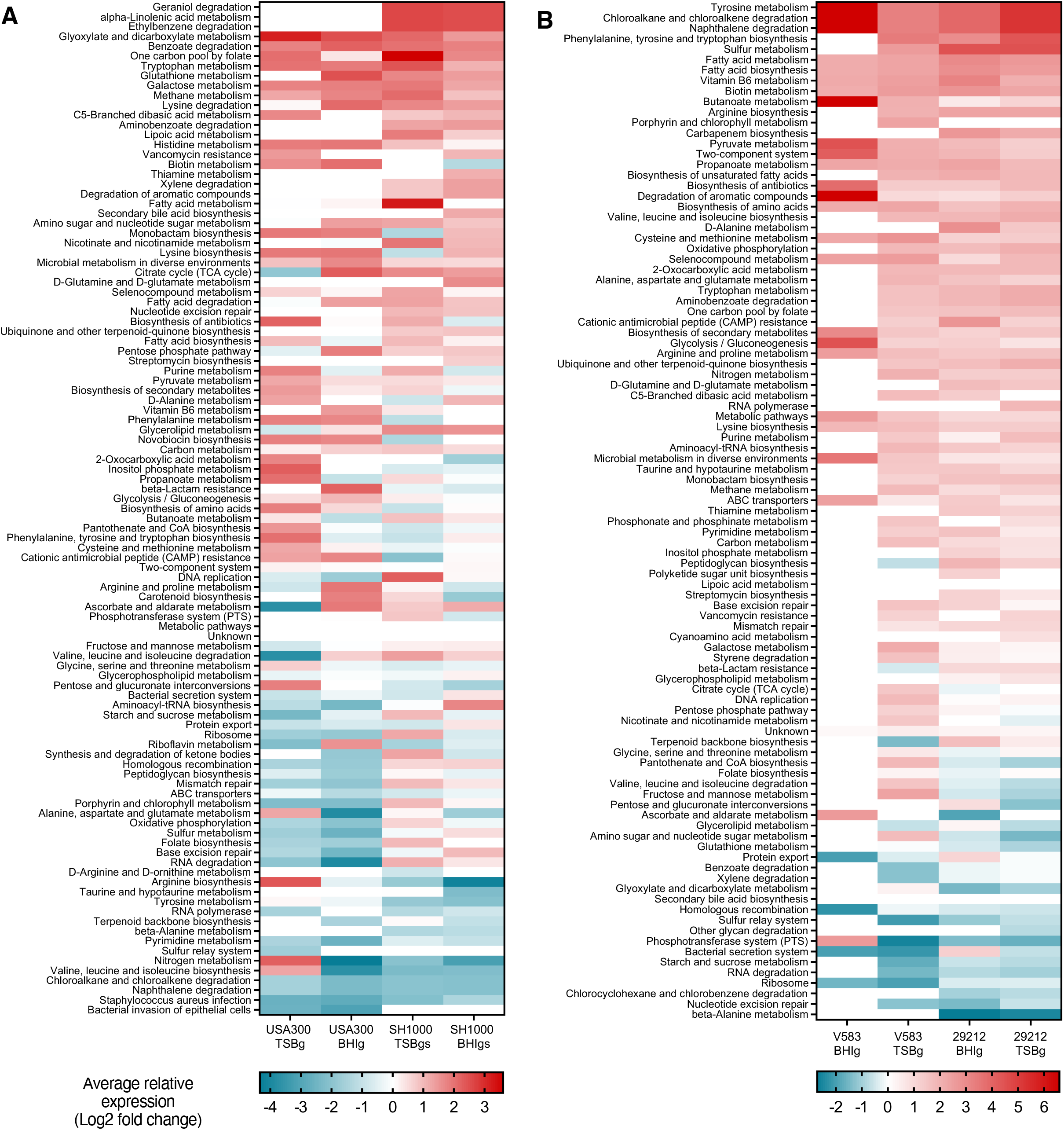
Relative expression of KEGG terms during biofilm formation. Each row of both heatmaps represents a single functional annotation term from DAVID Bioinformatic software using KEGG pathways annotation source. The term’s relative expression is the average of the log2 fold change of every associated gene with an adjusted *P*-value ≤ 0.05 from the transcriptomic analysis. The red (upregulated) and blue (downregulated) colour scale indicates relative expression level in average log2 fold change. (A) Functional annotation terms’ (total = 103) relative expression in *S. aureus* USA300 and SH1000, in BHIg(s) or TSBg(s) medium. (B) Functional annotation terms’ (total = 97) relative expression in *E. faecalis* V583 and 29212, in BHIg or TSBg medium.

**Figure S4.**
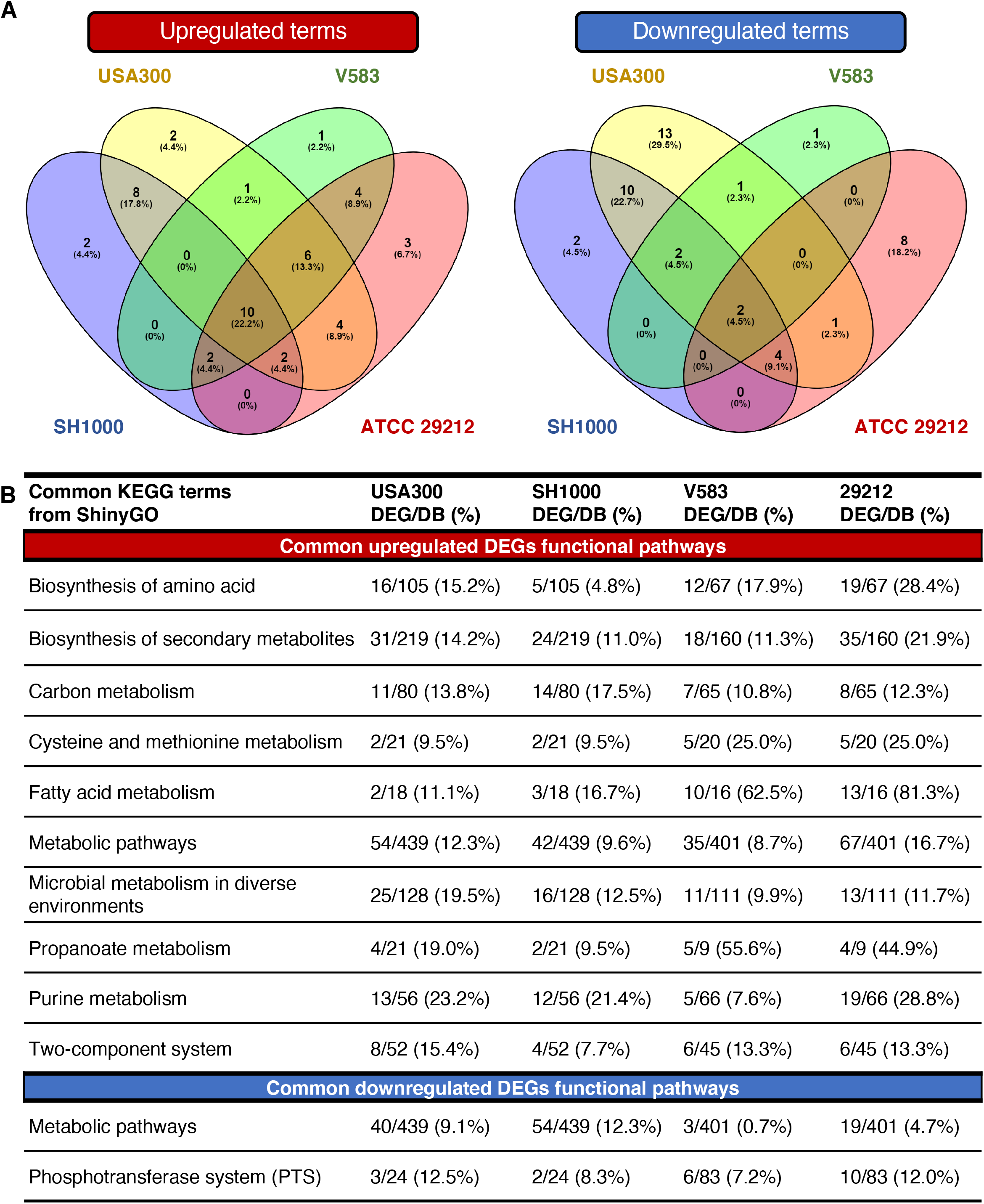
Common cellular adaptation between *S. aureus* and *E. faecalis* strains according to ShinyGO’s database. (A) Venn diagram representation of the unique and common functional terms of the upregulated DEGs (*P*-value ≤ 0.05, log2 fold change ≥ 2.0) and downregulated DEGs (*P*-value ≤ 0.05, log2 fold change ≤ −2.0) between *S. aureus* USA300 (yellow), *S. aureus* SH1000 (blue), *E. faecalis* V583 (green) and *E. faecalis* 29212 (red). (B) Table of the common cellular pathways between the four strains of this study, either upregulated or downregulated. GSEA was performed with ShinyGO using KEGG pathways and GO:BP annotation source. DEG/DB (%) is the number of DEGs submitted to ShinyGO reported on the number of genes existing in the database for the corresponding functional term (DB).

## References

1. Flemming HC, Wuertz S. 2019. Bacteria and archaea on Earth and their abundance in biofilms. Nat Rev Microbiol 17:247–260.

2. Flemming H-C, Wingender J, Szewzyk U, Steinberg P, Rice SA, Kjelleberg S. 2016. Biofilms: an emergent form of bacterial life. Nat Rev Microbiol 14:563–575.

3. Donlan RM. 2002. Biofilms: Microbial life on surfaces. Emerg Infect Dis. 8(9):881–90

4. Galié S, García-Gutiérrez C, Miguélez EM, Villar CJ, Lombó F. 2018. Biofilms in the food industry: Health aspects and control methods. Front Microbiol. May 7, 2018

5. Sahoo D, Bhatt M, Jena S, Dash D, Chayani N. 2015. Study of biofilm in bacteria from water pipelines. J Clin Diagnostic Res 9:9–11.

6. Høiby N, Ciofu O, Johansen HK, Song ZJ, Moser C, Jensen PØ, Molin S, Givskov M, Tolker-Nielsen T, Bjarnsholt T. 2011. The clinical impact of bacterial biofilms, *In* International Journal of Oral Science. p. 55–65.

7. Costerton JW, Cheng KJ, Geesey GG, Ladd TI, Nickel JC, Dasgupta M, Marrie TJ. 1987. Bacterial Biofilms in Nature and Disease. Annu Rev Microbiol 41:435–464.

8. Jamal M, Ahmad W, Andleeb S, Jalil F, Imran M, Nawaz MA, Hussain T, Ali M, Rafiq M, Kamil MA. 2018. Bacterial biofilm and associated infections. J Chinese Med Assoc. 81(1):7–11

9. Van Epps JS, Younger JG. 2016. Implantable device-related infection. Shock 46:597–608.

10. Lewis K. 2001. Riddle of biofilm resistance. Antimicrob Agents Chemother. 45(4):999–1007.

11. Arciola CR, Campoccia D, Montanaro L. 2018. Implant infections: adhesion, biofilm formation and immune evasion. Nat Rev Microbiol 16:1–13.

12. Public Health Agency of Canada. 2014. Central Venous Catheter-Associated Blood Stream Infections in Intensive Care Units in Canadian Acute-Care Hospitals - Surveillance Report. Centre for Communicable Diseases and Infection Control, Public Health Agency of Canada. https://www.ammi.ca/Guideline/24.ENG.pdf

13. Davies D. 2003. Understanding biofilm resistance to antibacterial agents. Nat Rev Drug Discov 2:114–22.

14. Høiby N, Bjarnsholt T, Givskov M, Molin S, Ciofu O. 2010. Antibiotic resistance of bacterial biofilms. Int J Antimicrob Agents 35:322–32.

15. Khatoon Z, McTiernan CD, Suuronen EJ, Mah TF, Alarcon EI. 2018. Bacterial biofilm formation on implantable devices and approaches to its treatment and prevention. Heliyon. 4(12):e01067

16. Lebeaux D, Ghigo J-M, Beloin C. 2014. Biofilm-Related Infections: Bridging the Gap between Clinical Management and Fundamental Aspects of Recalcitrance toward Antibiotics. Microbiol Mol Biol Rev 78:510–543.

17. Justo JA, Bookstaver PB. 2014. Antibiotic lock therapy: Review of technique and logistical challenges. Infect Drug Resist. 12(7):343–63

18. Kluytmans J, Van Belkum A, Verbrugh H. 1997. Nasal carriage of *Staphylococcus aureus*: Epidemiology, underlying mechanisms, and associated risks. Clin Microbiol Rev. 3(10):505–20

19. Mack D, Becker P, Chatterjee I, Dobinsky S, Knobloch JKM, Peters G, Rohde H, Herrmann M. 2004. Mechanisms of biofilm formation in *Staphylococcus epidermidis* and *Staphylococcus aureus*: Functional molecules, regulatory circuits, and adaptive responses. Int J Med Microbiol. 294(2-3):203–12

20. Beenken KE, Dunman PM, McAleese F, Macapagal D, Murphy E, Projan SJ, Blevins JS, Smeltzer MS. 2004. Global gene expression in *Staphylococcus aureus* biofilms. J Bacteriol 186:4665–4684.

21. Cue D, Lei MG, Lee CY. 2012. Genetic regulation of the intercellular adhesion locus in staphylococci. Front Cell Infect Microbiol. 2:38

22. Fitzpatrick F, Humphreys H, O’Gara JP. 2005. Evidence for icaADBC-independent biofilm development mechanism in methicillin-resistant *Staphylococcus aureus* clinical isolates. J Clin Microbiol 43:1973–1976.

23. Otto M. 2013. Staphylococcal Infections: Mechanisms of Biofilm Maturation and Detachment as Critical Determinants of Pathogenicity. Annu Rev Med 64:175–188.

24. O’Gara JP. 2007. ica and beyond: Biofilm mechanisms and regulation in *Staphylococcus epidermidis* and *Staphylococcus aureus*. FEMS Microbiol Lett. 270(2):179–88

25. McCarthy H, Rudkin JK, Black NS, Gallagher L, O’Neill E, O’Gara JP. 2015. ethicillin resistance and the biofilm phenotype in *Staphylococcus aureus*. Front Cell Infect Microbiol 5:1–9.

26. Mlynek KD, Bulock LL, Stone CJ, Curran LJ, Marat R, Bayles KW, Brinsmade SR. 2020. Genetic and biochemical analysis of CodY-mediated cell aggregation in *Staphylococcus aureus* reveals an interaction between eDNA and polysaccharide in the extracellular matrix. J Bacteriol 202(8):e00593–19

27. Willett JLE, Ji MM, Dunny GM. 2019. Exploiting biofilm phenotypes for functional characterization of hypothetical genes in *Enterococcus faecalis*. npj Biofilms Microbiomes 5:1–14.

28. Edmond MB, Ober JF, Dawson JD, Weinbaum DL, Wenzel RP. 1996. Vancomycin-Resistant Enterococcal Bacteremia: Natural History and Attributable Mortality. Clin Infect Dis 23:1234–1239.

29. Fisher K, Phillips C. 2009. The ecology, epidemiology and virulence of Enterococcus. Microbiology. 155(Pt 6):1749–1757

30. Flahaut S, Hartke A, Giard J, Benachour A, Boutibonnes P, Auffray Y. 1996. Relationship between stress response towards bile salts, acid and heat treatment in *Enterococcus faecalis*. FEMS Microbiol Lett 138:49–54.

31. Huycke MM, Sahm DF, Gilmore MS. 1998. Multiple-drug resistant enterococci: The nature of the problem and an agenda for the future. Emerg Infect Dis. 4(2):239–49

32. Wang X, He X, Jiang Z, Wang J, Chen X, Liu D, Wang F, Guo Y, Zhao J, Liu F, Huang L, Yuan J. 2010. Proteomic analysis of the *Enterococcus faecalis* V583 strain and clinical isolate V309 under vancomycin treatment. J Proteome Res 9:1772–1785.

33. Yin W, Wang Y, Liu L, He J. 2019. Biofilms: The microbial “protective clothing” in extreme environments. Int J Mol Sci 20(14):3423

34. Ch’ng JH, Chong KKL, Lam LN, Wong JJ, Kline KA. 2019. Biofilm-associated infection by enterococci. Nat Rev Microbiol. 17(2):82–94

35. Pisithkul T, Schroeder JW, Trujillo EA, Yeesin P, Stevenson DM, Chaiamarit T, Coon JJ, Wang JD, Amador-Noguez D. 2019. Metabolic Remodeling during Biofilm Development of *Bacillus subtilis*. MBio 10:1–32.

36. Caro-Astorga J, Frenzel E, Perkins JR, Álvarez-Mena A, de Vicente A, Ranea JAG, Kuipers OP, Romero D. 2020. Biofilm formation displays intrinsic offensive and defensive features of *Bacillus cereus*. npj Biofilms Microbiomes 6:3.

37. Zhu Y, Xiong YQ, Sadykov MR, Fey PD, Lei MG, Lee CY, Bayer AS, Somerville GA. 2009. Tricarboxylic acid cycle-dependent attenuation of *Staphylococcus aureus in vivo* virulence by selective inhibition of amino acid transport. Infect Immun 77:4256–4264.

38. Moormeier DE, Bayles KW. 2017. *Staphylococcus aureus* biofilm: a complex developmental organism. Mol Microbiol 104:365–376.

39. Tendolkar PM, Baghdayan AS, Shankar N. 2006. Putative surface proteins encoded within a novel transferable locus confer a high-biofilm phenotype to *Enterococcus faecalis*. J Bacteriol 188:2063–72.

40. Zhu Y, Weiss EC, Otto M, Fey PD, Smeltzer MS, Somerville GA. 2007. *Staphylococcus aureus* biofilm metabolism and the influence of arginine on polysaccharide intercellular adhesin synthesis, biofilm formation, and pathogenesis. Infect Immun 75:4219–4226.

41. Moormeier DE, Bose JL, Horswill AR, Bayles KW. 2014. Temporal and stochastic control of *Staphylococcus aureus* biofilm development. MBio 5(5):e01341–14.

42. Wattam AR, Davis JJ, Assaf R, Boisvert S, Brettin T, Bun C, Conrad N, Dietrich EM, Disz T, Gabbard JL, Gerdes S, Henry CS, Kenyon RW, Machi D, Mao C, Nordberg EK, Olsen GJ, Murphy-Olson DE, Olson R, Overbeek R, Parrello B, Pusch GD, Shukla M, Vonstein V, Warren A, Xia F, Yoo H, Stevens RL. 2016. Improvements to PATRIC, the all-bacterial Bioinformatics Database and Analysis Resource Center. Nucleic Acids Res 45:535–542.

43. Yan F, Yu Y, Gozzi K, Chen Y, Guo J, Chai Y. 2017. Genome-Wide Investigation of Biofilm Formation in *Bacillus cereus*. Appl Environ Microbiol 83:e00561–17.

44. Sadovskaya I, Vinogradov E, Flahaut S, Kogan G, Jabbouri S. 2005. Extracellular carbohydrate-containing polymers of a model biofilm-produring strain, *Staphylococcus epidermidis* RP62A. Infect Immun 73:3007–3017.

45. Sugimoto S, Sato F, Miyakawa R, Chiba A, Onodera S, Hori S, Mizunoe Y. 2018. Broad impact of extracellular DNA on biofilm formation by clinically isolated Methicillin-resistant and -sensitive strains of *Staphylococcus aureus*. Sci Rep 8(1):2254

46. Mohamed JA, Huang DB. 2007. Biofilm formation by enterococci. J Med Microbiol 56:1581–1588.

47. Huang DW, Sherman BT, Lempicki RA. 2009. Systematic and integrative analysis of large gene lists using DAVID bioinformatics resources. Nat Protoc 4:44–57.

48. Karp PD, Paley SM, Midford PE, Krummenacker M, Billington R, Kothari A, Ong WK, Subhraveti P, Keseler IM, Caspi R. 2015. Pathway Tools version 23.0: Integrated Software for Pathway/Genome Informatics and Systems Biology. Brief Bioinform 11:40–79.

49. Fuchs S, Mehlan H, Bernhardt J, Hennig A, Michalik S, Surmann K, Pané-Farré J, Giese A, Weiss S, Backert L, Herbig A, Nieselt K, Hecker M, Völker U, Mäder U. 2018. AureoWiki ̵ The repository of the *Staphylococcus aureus* research and annotation community. Int J Med Microbiol 308:558–568.

50. Ge SX, Jung D, Jung D, Yao R. 2020. ShinyGO: A graphical gene-set enrichment tool for animals and plants. Bioinformatics 36:2628–2629.

51. Kilstrup M, Hammer K, Ruhdal Jensen P, Martinussen J. 2005. Nucleotide metabolism and its control in lactic acid bacteria. FEMS Microbiol Rev 29:555–590.

52. Bose JL, Fey PD, Bayles KW. 2013. Genetic tools to enhance the study of gene function and regulation in *Staphylococcus aureus*. Appl Environ Microbiol 79:2218–2224.

53. Zhang Y, Morar M, Ealick SE. 2008. Structural biology of the purine biosynthetic pathway. Cell Mol Life Sci 65:3699–3724.

54. O’Toole GA. 2011. Microtiter dish Biofilm formation assay. J Vis Exp 47:2437

55. Langmead B, Salzberg SL. 2012. Fast gapped-read alignment with Bowtie 2. Nat Methods 9:357–9.

56. Herbert S, Ziebandt AK, Ohlsen K, Schäfer T, Hecker M, Albrecht D, Novick R, Götz F. 2010. Repair of global regulators in Staphylococcus aureus 8325 and comparative analysis with other clinical isolates. Infect Immun 78:2877–2889.

57. Love MI, Huber W, Anders S. 2014. Moderated estimation of fold change and dispersion for RNA-seq data with DESeq2. Genome Biol 15:550.

58. Olivieros JC. 2007. VENNY. An interactive tool for comparing lists with Venn Diagrams. https://bioinfogp.cnb.csic.es/tools/venny_old/venny.php

59. Malheiro J, Simões M. 2016. Antimicrobial resistance of biofilms in medical devices, p. 98–113. *In* Biofilms and Implantable Medical Devices: Infection and Control. Elsevier Inc.

60. Floyd KA, Eberly AR, Hadjifrangiskou M. 2017. Adhesion of bacteria to surfaces and biofilm formation on medical devices, p. 47–95. *In* Biofilms and Implantable Medical Devices: Infection and Control. Elsevier Inc.

61. Tendolkar PM, Baghdayan AS, Gilmore MS, Shankar N. 2004. Enterococcal surface protein, Esp, enhances biofilm formation by *Enterococcus faecalis*. Infect Immun 72:6032–9.

62. Dogsa I, Brloznik M, Stopar D, Mandic-Mulec I. 2013. Exopolymer Diversity and the Role of Levan in *Bacillus subtilis* Biofilms. PLoS One 8:2–11.

63. Stipetic LH, Dalby MJ, Davies RL, Morton FR, Ramage G, Burgess KEV. 2016. A novel metabolomic approach used for the comparison of *Staphylococcus aureus* planktonic cells and biofilm samples. Metabolomics 12:75.

64. Ammons MCB, Tripet BP, Carlson RP, Kirker KR, Gross MA, Stanisich JJ, Copié V. 2014. Quantitative NMR metabolite profiling of methicillin-resistant and methicillin-susceptible *Staphylococcus aureus* discriminates between biofilm and planktonic phenotypes. J Proteome Res 13:2973–85.

65. Resch A, Leicht S, Saric M, Pásztor L, Jakob A, Götz F, Nordheim A. 2006. Comparative proteome analysis of *Staphylococcus aureus* biofilm and planktonic cells and correlation with transcriptome profiling. Proteomics 6:1867–1877.

66. Resch A, Rosenstein R, Nerz C, Götz F. 2005. Differential gene expression profiling of *Staphylococcus aureus* cultivated under biofilm and planktonic conditions. Appl Environ Microbiol 71:2663–2676.

67. Liu J, Hou Y, Peters BM, Su J, Li L, Li B, Chen D, Li Y, Xu Z, Shirtliff ME. 2018. Transcriptomics study on *Staphylococcus aureus* biofilm under low concentration of ampicillin. Front Microbiol 9:2413.

68. Seneviratne CJ, Suriyanarayanan T, Swarup S, Chia KHB, Nagarajan N, Zhang C. 2017. Transcriptomics Analysis Reveals Putative Genes Involved in Biofilm Formation and Biofilm-associated Drug Resistance of *Enterococcus faecalis*. J Endod 43:949–955.

69. Suryaletha K, Narendrakumar L, John J, Radhakrishnan MP, George S, Thomas S. 2019. Decoding the proteomic changes involved in the biofilm formation of *Enterococcus faecalis* SK460 to elucidate potential biofilm determinants. BMC Microbiol 19:146.

70. Philips J, Rabaey K, Lovley DR, Vargas M. 2017. Biofilm formation by *Clostridium ljungdahlii* is induced by sodium chloride stress: Experimental evaluation and transcriptome analysis. PLoS One 12:1–25.

71. Kim JK, Kwon JY, Kim SK, Han SH, Won YJ, Lee JH, Kim CH, Fukatsu T, Lee BL. 2014. Purine biosynthesis, biofilm formation, and persistence of an insect-microbe gut symbiosis. Appl Environ Microbiol 80:4374–4382.

72. Shaffer CL, Zhang EW, Dudley AG, Dixon BREA, Guckes KR, Breland EJ, Floyd KA, Casella DP, Algood HMS, Clayton DB, Hadjifrangiskou M. 2017. Purine biosynthesis metabolically constrains intracellular survival of uropathogenic *Escherichia coli*. Infect Immun 85(1):e00471–16

73. Yoshioka S, Newell PD. 2016. Disruption of de novo purine biosynthesis in *Pseudomonas fluorescens* Pf0-1 leads to reduced biofilm formation and a reduction in cell size of surface-attached but not planktonic cells. PeerJ 2016:1–24.

74. Ge X, Kitten T, Chen Z, Lee SP, Munro CL, Xu P. 2008. Identification of *Streptococcus sanguinis* genes required for biofilm formation and examination of their role in endocarditis virulence. Infect Immun 76:2551–2559.

75. Dengler V, Foulston L, DeFrancesco AS, Losick R. 2015. An electrostatic net model for the role of extracellular DNA in biofilm formation by *Staphylococcus aureus*. J Bacteriol 197:3779–3787.

76. Thomas VC, Thurlow LR, Boyle D, Hancock LE. 2008. Regulation of autolysis-dependent extracellular DNA release by *Enterococcus faecalis* extracellular proteases influences biofilm development. J Bacteriol 190:5690–5698.

77. Vilain S, Pretorius JM, Theron J, Brözel VS. 2009. DNA as an adhesin: *Bacillus cereus* requires extracellular DNA to form biofilms. Appl Environ Microbiol 75:2861–2868.

78. Fahmi T, Faozia S, Port GC, Cho KH. 2019. The second messenger C-di-AMP regulates diverse cellular pathways involved in stress response, biofilm formation, cell wall homeostasis, speb expression, and virulence in streptococcus pyogenes. Infect Immun 87:1–19.

79. Chávez de Paz LE, Lemos JA, Wickström C, Sedgley CM. 2012. Role of (p)ppGpp in biofilm formation by *Enterococcus faecalis*. Appl Environ Microbiol 78:1627–30.

